# Deciphering the functioning of microbial communities: shedding light on the critical steps in metaproteomics

**DOI:** 10.1101/697599

**Authors:** Augustin Géron, Johannes Werner, Ruddy Wattiez, Philippe Lebaron, Sabine Matallana-Surget

**Affiliations:** University of Stirling, Division of Biological and Environmental Sciences, Faculty of Natural Sciences, Stirling, UK; University of Mons, Proteomic and Microbiology Department, Mons, Belgium; Leibniz Institute for Baltic Sea Research, Department of Biological Oceanography, Rostock-Warnemünde, Germany; Sorbonne Universités, UPMC Univ Paris 06, USR3579, LBBM, Observatoire Océanologique, 66651 Banyuls/mer, France

**Keywords:** Metaproteomics, Metagenomics, Bioinformatics, Mass Spectrometry, Microbial Ecology

## Abstract

Unraveling the complex structure and functioning of microbial communities is essential to accurately predict the impact of perturbations and/or environmental changes. From all molecular tools available today to resolve the dynamics of microbial communities, metaproteomics stands out, allowing the establishment of phenotype-genotype linkages. Despite its rapid development, this technology has faced many technical challenges that still hamper its potential power. How to maximize the number of protein identification, improve quality of protein annotation and provide reliable ecological interpretation, are questions of immediate urgency. In our study, we used a robust metaproteomic workflow combining two protein fractionation approaches (gel-based *versus* gel-free) and four protein search databases derived from the same metagenome to analyze the same seawater sample. The resulting eight metaproteomes provided different outcomes in terms of (i) total protein numbers, (ii) taxonomic structures, and (iii) protein functions. The characterization and/or representativeness of numerous proteins from ecologically relevant taxa such as *Pelagibacterales*, *Rhodobacterales* and *Synechococcales*, as well as crucial environmental processes, such as nutrient uptake, nitrogen assimilation, light harvesting and oxidative stress response were found to be particularly affected by the methodology. Our results provide clear evidences that the use of different protein search databases significantly alters the biological conclusions in both gel-free and gel-based approaches. Our findings emphasize the importance of diversifying the experimental workflow for a comprehensive metaproteomic study.

## Background

Metaproteomics aims at characterizing the total proteins obtained from microbial communities [1] and, in association with metagenomics, unraveling the functional complexity of a given ecosystem [2]. Since the first environmental metaproteomic study performed in the Chesapeake Bay [3], numerous investigations were carried out in a variety of environments using descriptive, comparative and/or quantitative approaches [4]. Comparative metaproteomics was often used to describe spatial and seasonal changes in aquatic ecosystems using (i) *in situ* [5–8] (ii) mesocosms [9, 10] or (iii) microcosms [11] approaches.

Metaproteomics on marine ecosystems is a rapidly expanding field that involves a series of challenging steps and critical decisions in its workflow [4, 12–14]. The marine metaproteomic workflow consists mainly of four steps: (i) sampling and protein extraction, (ii) protein separation, (iii) mass spectrometry, and (iv) protein identification/annotation [15]. Until now, standardized experimental protocols are still missing, leading to methodological inconsistencies and data interpretation biases across metaproteomic studies [16–18].

Protein identification strongly relies on both the quality of experimental mass spectra and the comprehensiveness of the protein search database (DB) [15]. Both gel-based [19] and shotgun gel-free [5, 20] approaches have been used in metaproteomic analyses and both were found to be complementary [4]. Two main data sources are commonly used to construct protein search DB: public protein repositories, and/or metagenomic data [13]. Identifying proteins by searching against public protein repositories such as UniProtKB/SwissProt, UniProtKB/TrEMBL, UniRef, NCBI, or Ensembl is challenging because of the large size of these DBs, which increase search space and overestimate false discovery rate (FDR), thus decreasing the total number of identified proteins [17, 18, 21, 22]. To address the issue of large size DB, different strategies were developed such as taxonomy-based filtration [13], partial searches against smaller sub-DB [23, 24] or the two-round DB searching method [22]. The two-round DB searching method consists in searching experimental mass spectra against a refined database composed of the protein sequences identified in a preliminary error tolerant search, allowing significant increase in the total number of identified proteins. This strategy was extensively used in recent metaproteomics studies [25–28]. Regarding metagenomic data, both assembled [6] and non-assembled [24, 30] sequencing reads were used in metaproteomics for protein search DB creation. Skipping read assembly was shown to prevent information loss and potential noise introduction and led to higher protein identification yield [29].

Metaproteomic data analysis also involves taxonomic and functional annotation. Due to the protein inference issue (i.e. a same peptide can be found in homologous proteins), inaccurate protein annotations are commonly encountered in metaproteomics [30]. To overcome this issue, protein identification tools such as Pro Group algorithm (Protein Pilot) [31], Prophane [32] or MetaProteomeAnalyzer [33] automatically group homologous protein sequences. In our study, we used mPies program [34], which computes taxonomic consensus annotation on protein groups using LCA [13, 35] and provides a novel consensus functional annotation based on Uniprot DB giving more accurate insights into the diversity of protein functions compared to former strategies mapping proteins on broader functional categories, such as KEGG [36] or COGs [37].

To what extend the methodology affects the metaproteome interpretation has already been studied in artificial microbial communities [17] and gut microbiomes [24, 38] but its impact on marine samples still remains poorly documented [18]. In this study, we used a robust experimental design comparing the combined effect of protein search DB choice and protein fractionation approach on the same sea surface sample. For this purpose, two sets of peptide spectra resulting from gel-based and gel-free approaches were searched against four DBs derived from the same raw metagenomic data. The resulting eight metaproteomes were quantitatively and qualitatively compared, demonstrating to which extent diversifying metaproteomic workflow allows the most comprehensive understanding of microbial communities dynamics.

## Materials and methods

### Sampling

Seawater samples (n=4) were collected in summer (June 2014) at the SOLA station, located 500 m offshore of Banyuls-sur-mer, in the Northwestern Mediterranean Sea (42° 49’ N, 3° 15’ W). Each sample consisted of 60 liters of sea surface water, pre-filtered at 5 µm and subsequently sequentially filtered through 0.8 and 0.2 µm pore-sized filters (polyethersulfone membrane filters, PES, 142 mm, Millipore). Four independent sets of filters were obtained and flash frozen into liquid nitrogen before storage at – 80 °C.

### Protein isolation for gel-based and gel-free approaches

A combination of different physical (sonication/freeze–thaw) and chemical (urea/thiourea containing buffers, acetone precipitation) extraction techniques were used on the filtered seawater samples to maximize the recovery of protein extracts from the filters. The 0.2 µm filters were removed from their storage buffer and cut into quarters using aseptic procedures. Protein isolation was performed on four 0.2 µm filters. The same protein isolation protocol was used for both gel-based and gel-free approaches. The filters were suspended in a lysis buffer containing 8 M Urea / 2 M Thiourea, 10 mM HEPES, and 10 mM dithiothreitol. Filters were subjected to five freeze–thaw cycles in liquid N2 to release cells from the membrane. Cells were mechanically broken by sonication on ice (5 cycles of 1 min with tubes on ice, amplitude 40 %, 0.5 pulse rate) and subsequently centrifuged at 16 000 g at 4 °C for 15 min. To remove particles that did not pellet during the centrifugation step, we filtered the protein suspension through a 0.22 mm syringe filter and transferred into a 3 kDa cutoff Amicon Ultra-15 filter unit (Millipore) for protein concentration. Proteins were precipitated with cold acetone overnight at −80 °C, with an acetone/aqueous protein solution ratio of 4:1. Total protein concentration was determined by a Bradford assay, according to the Bio-Rad Protein Assay kit (Bio-Rad, Hertfordshire, UK) according to manufacturer’s instructions, with bovine γ-globulin as a protein standard. Protein samples were reduced with 25 mM dithiothreitol (DTT) at 56 °C for 30 min and alkylated with 50 mM iodoacetamide at room temperature for 30 min. For gel-free liquid chromatography tandem mass spectrometry analysis, a trypsic digestion (sequencing grade modified trypsin, Promega) was performed overnight at 37 °C, with an enzyme/substrate ratio of 1:25.

### Gel-based proteomics approach

Protein isolates diluted in Laemmli buffer (2 % SDS, 10 % glycerol, 5 % β-mercaptoethanol, 0.002 % bromophenol blue and 0.125 M Tris-HCl, pH 6.8) and sonicated in a water bath six times for 1 min at room temperature. After 1 min incubation at 90 °C, the protein solutions were centrifuged at 13 000 rpm at room temperature for 15 min. The SDS-PAGE of the protein mixtures was conducted using 4–20 % precast polyacrylamide mini-gels (Pierce). The protein bands were visualized with staining using the Imperial Protein Stain (Thermo) according to the manufacturer’s instructions. The corresponding gel lane containing proteins was cut in 17 pieces of 1 mm each. Enzymatic digestion was performed by the addition of 10 µL modified sequencing grade trypsin (0.02 mg/mL) in 25 mM NH_4_HCO_3_ to each gel piece. The samples were placed for 15 min at 4 °C and incubated overnight at 37 °C. The reaction was stopped with 1 µL 5 % (v/v) formic acid. Tryptic peptides were analyzed by liquid chromatography tandem mass spectrometry.

### Liquid chromatography tandem mass spectrometry analysis

Purified peptides from digested protein samples from gel-free and gel-based proteomics were identified using a label-free strategy on an UHPLC-HRMS platform composed of an eksigent 2D liquid chromatograph and an AB SCIEX Triple TOF 5 600. Peptides were separated on a 25 cm C18 column (Acclaim pepmap 100, 3 μm, Dionex) by a linear acetonitrile (ACN) gradient [5–35 % (v/v), in 15 or 120 min] in water containing 0.1 % (v/v) formic acid at a flow rate of 300 nL min-1. Mass spectra (MS) were acquired across 400–1,500 m/z in high-resolution mode (resolution > 35 000) with 500 ms accumulation time. Six microliters of each fraction were loaded onto a pre-column (C18 Trap, 300 µm i.d.×5 mm, Dionex) using the Ultimate 3000 system delivering a flow rate of 20 µL/min loading solvent (5 % (v/v) acetonitrile (ACN), 0.025 % (v/v) TFA). After a 10 min desalting step, the pre-column was switched online with the analytical column (75 µm i.d.×15 cm PepMap C18, Dionex) equilibrated in 96 % solvent A (0.1 % (v/v) formic acid in HPLC-grade water) and 4 % solvent B (80 % (v/v) ACN, 0.1 % (v/v) formic acid in HPLC-grade water). Peptides were eluted from the pre-column to the analytical column and then to the mass spectrometer with a gradient from 4-57 % solvent B for 50 min and 57-90 % solvent B for 10 min at a flow rate of 0.2 µL min−1 delivered by the Ultimate pump. Positive ions were generated by electrospray and the instrument was operated in a data-dependent acquisition mode described as follows: MS scan range: 300 – 1 500 m/z, maximum accumulation time: 200 ms, ICC target: 200 000. The top 4 most intense ions in the MS scan were selected for MS/MS in dynamic exclusion mode: ultrascan, absolute threshold: 75 000, relative threshold: 1 %, excluded after spectrum count: 1, exclusion duration: 0.3 min, averaged spectra: 5, and ICC target: 200 000. Gel-based and gel-free metaproteomic data were submitted to iProx [39] (Project ID: IPX0001684000).

### Databases creation and protein identification

Protein searches were performed with ProteinPilot (ProteinPilot Software 5.0.1; Revision: 4895; Paragon Algorithm: 5.0.1.0.4874; AB SCIEX, Framingham, MA) (Matrix Science, London, UK; v. 2.2). Paragon searches 34 were conducted using LC MS/MS Triple TOF 5600 System instrument settings. Other parameters used for the search were as follows: Sample Type: Identification, Cys alkylation: Iodoacetamide, Digestion: Trypsin, ID Focus: Biological Modifications and Amino acid substitutions, Search effort: Thorough ID, Detected Protein Threshold [Unused ProtScore (Conf)] >: 0.05 (10.0%).

Three DBs were created using the same metagenome (Project number: ERP009703, Ocean Sampling Day 2014, sample: OSD14_2014_06_2m_NPL022, run ID: ERR771073) and were generated with mPies v 0.9, our recently in house developed mPies program freely available at https://github.com/johanneswerner/mPies/ (Additional file 4) [34]. The three DBs were: (i) a non-assembled metagenome-derived DB (NAM-DB), (ii) an assembled metagenome-derived DB (AM-DB) and (iii) a taxonomy-derived DB (TAX-DB) (Table 1). Briefly, mPies first trimmed sequencing raw reads with Trimmomatic [40]. For NAM-DB, mPies directly predicted genes from trimmed sequencing reads with FragGeneScan [41]. For AM-DB, mPies first assembled trimmed sequencing reads into contigs using metaSPAdes [42] and subsequently called genes with Prodigal [43]. For TAX-DB, mPies used SingleM [44] to predict operational taxonomic units from the trimmed sequencing reads and retrieved all the taxon IDs at genus level. All available proteomes for each taxon ID were subsequently downloaded from UniProtKB/TrEMBL. Duplicated protein sequences were removed with CD-HIT [45] from each DB.

**Table 1.** Two-round search performances obtained for each methodology. Searching parameters are provided in the *Material and methods* section.

|  | Database | Number of proteins in database | Number of peptide spectra identified <sup>a</sup> | Coverage of peptide spectra identified (%) | Number of distinct peptides Identified <sup>a</sup> | Number of proteins after validation <sup>a,b</sup> |
| --- | --- | --- | --- | --- | --- | --- |
| *First-round searches* |  |  |  |  |  |  |
| **Gel-free** | AM-DB | 64 613 | 24 684 | 7.8 | 3 237 | 347 |
|  | NAM-DB | 462 821 | 35 430 | 11.1 | 4 295 | 834 |
|  | TAX-DB | 13 426 277 | 10 626 | 3.3 | 1 624 | 607 |
| **Gel-based** | AM-DB | 64 613 | 2 066 | 24.7 | 1 408 | 201 |
|  | NAM-DB | 462 821 | 2 584 | 30.9 | 1 849 | 652 |
|  | TAX-DB | 13 426 277 | 2 304 | 27.5 | 1 526 | 496 |
| *Second-round searches* |  |  |  |  |  |  |
| **Gel-free** | AM-DB | 782 | 42 831 | 13.4 | 8 487 | 549 |
|  | NAM-DB | 4 277 | 57 840 | 18.2 | 9 113 | 1 131 |
|  | TAX-DB | 18 480 | 31 700 | 10.0 | 4 497 | 464 |
|  | Comb-DB | 23 405 | 56 530 | 17.7 | 8 273 | 1 048 |
| **Gel-based** | AM-DB | 377 | 2 619 | 31.3 | 1 815 | 277 |
|  | NAM-DB | 3 080 | 2 897 | 34.6 | 2 034 | 714 |
|  | TAX-DB | 19 036 | 2 951 | 35.3 | 1 777 | 434 |
|  | Comb-DB | 22 493 | 3 684 | 44.0 | 2 244 | 700 |

<sup>a</sup> values comprised in 95% confidence interval
<sup>b</sup> values at 1% global FDR and after manual validation for proteins identified with one peptide

Gel-based and gel-free MS/MS spectra were individually searched twice against the DBs. In the first-round search, full size NAM-DB, AM-DB and TAX-DB were used (Table 1). In the second-round search, each DB was restricted to the protein sequences identified in the first-round search. For both gel-free and gel-based approaches, the second round NAM-DB, AM-DB and TAX-DB were merged and redundant protein sequences were removed, leading to two combined DBs (Comb-DBs), subsequently searched against gel-based and gel-free MS/MS spectra. Consequently, a total of 8 metaproteomes obtained from four DBs: NAM-DB, AM-DB, TAX-DB and Comb-DB were analyzed in this paper. A FDR threshold of 1%, calculated at the protein level was used for each protein searches. Proteins identified with one single peptide were validated by manual inspection of the MS/MS spectra, ensuring that a series of at least five consecutive sequence-specific b-and y-type ions was observed.

### Protein annotation

Identified proteins were annotated using mPies. For taxonomic and functional annotation, mPies used Diamonds [46] to align each identified protein sequences against the non-redundant NCBI DB and the UniProt DB (Swiss-Prot) respectively and retrieved up to 20 best hits based on alignment score. For taxonomic annotation, mPies returned the last common ancestor (LCA) among the best hits via MEGAN (bit score >80) [35]. For functional annotation, mPies returned the most frequent protein name, with a consensus tolerance threshold above 80% of similarity amongst the 20 best blast hits. Proteins annotated with a score below this threshold were manually validated. Manual validation was straightforward as the main reasons leading to low annotation score were often explained by the characterization of protein isoforms or different sub-units of the same protein. To facilitate the understanding of this annotation step, examples were provided in Additional file 5. Annotated proteins files are available in Additional file 6.

## Results and discussion

### Database choice affects the total number of protein identification

The two-rounds search strategy commonly used in recent metaproteomics studies [25–28] significantly reduced the size of protein search DBs number, which in turn increased the total number of total identified proteins with both assembled metagenome-derived database (AM-DB) and non-assembled metagenome-derived database (NAM-DB) (Table 1). Overall, the total number of identified proteins was found to be consistent with other metaproteomics studies conducted in marine oligotrophic waters [7, 47–50]. NAM-DB led to greater protein identifications (gel-based: 714, gel-free: 1 131) than AM-DB (gel-based: 277 and gel-free: 549) and taxonomic-derived database (TAX-DB) (gel-based: 434 and gel-free: 464) for both proteomics approaches. Combined-database (Comb-DB) gave comparable results than NAM-DB in both approaches (gel-based: 700 and gel-free: 1 048). In AM-DB approach, the assembly process involved the removal of reads that cannot be assembled into longer contigs, leading to loss of gene fragments and consequently fewer identified proteins [51]. As high proportions of prokaryotic genomes are protein-coding, gene fragments can directly be predicted from non-assembled sequencing reads [52]. TAX-DB suffered from a reduction of protein detection sensitivity due to its large size, which negatively influenced FDR statistics and protein identification yield [22].

### Protein search DB affects the taxonomic structure

The proportion of proteins, for which a LCA was found, decreased with lowering taxonomic hierarchy (Domain > Phylum > Class > Order > Family > Genus), independently of the methodology (Figure 1). The proportion of annotated proteins at the domain, phylum and class levels remained constant with an average of 97.3 ± 1.0%, 92.0 ± 1.1% and 80.3 ± 0.8% respectively (Figure 1, Additional file 1). At order level and below, TAX-DB performed the best at assigning a LCA, in both gel-free and gel-based approaches. These results can be explained by the fact that the proteins were annotated using the program mPies which relies on protein sequence quality [34]. Indeed, TAX-DB comprised well-annotated protein sequences, obtained from bacterial proteomes retrieved from UniProtKB, while the other DBs used environmental reads, including fragmented and unsequenced bacterial genomes. This confirmed that the proportion of protein assigned to a LCA was affected by the DB choice, as previously demonstrated at the peptide level by May *et al*. (2016) [29].

**Figure 1.**
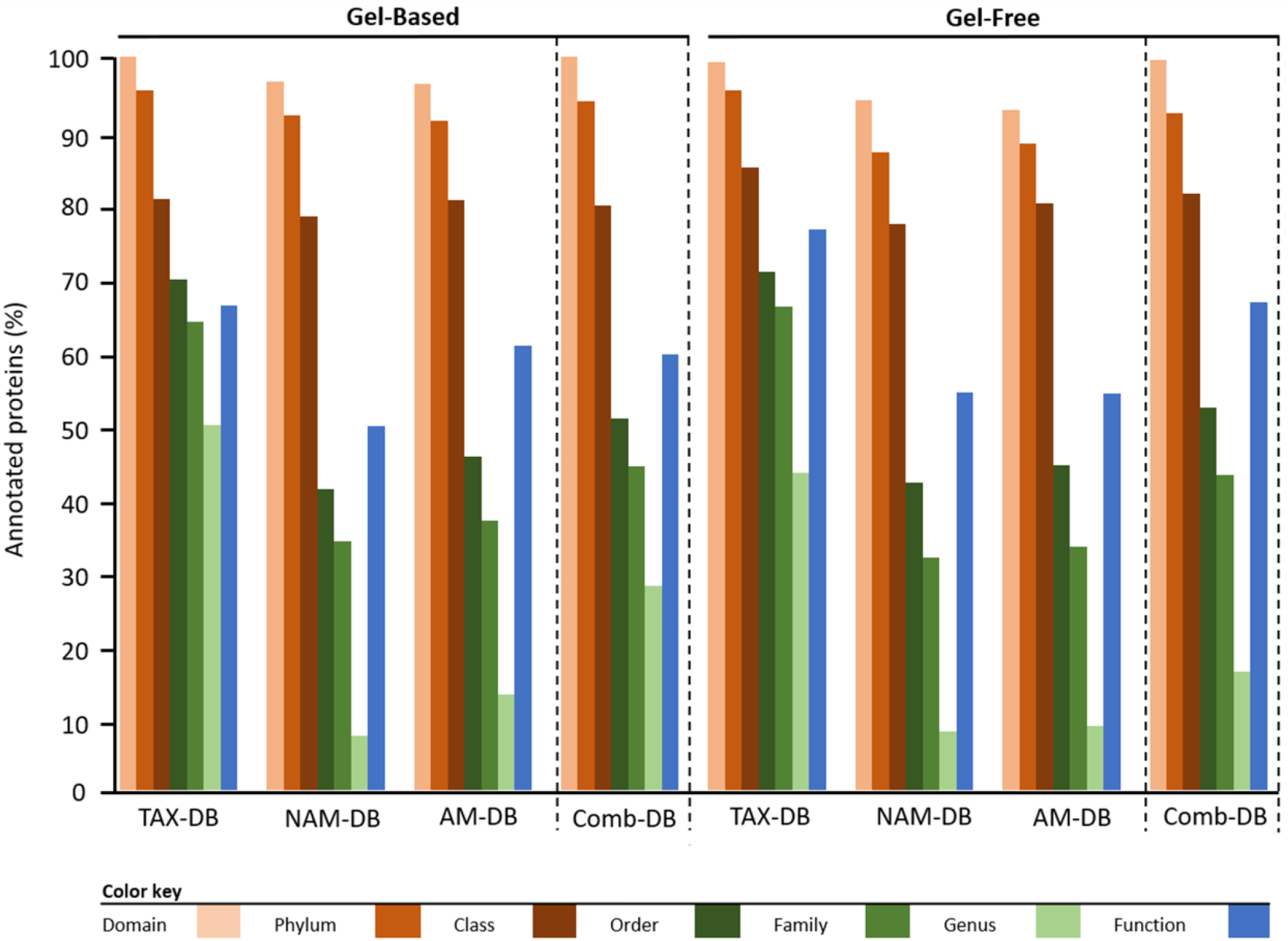
Taxonomic and functional protein annotation. Comparison of the proportion of proteins for which a consensus annotation was found. Bars represent the percentage of annotated proteins *versus* total identified proteins depending on methodology.

At phylum level, most of the proteins identified were assigned to *Proteobacteria* and the least abundant were mainly assigned to *Bacteroidetes* and *Cyanobacteria* (Table 2). Although *Proteobacteria* showed similar proportion in all metaproteomes (90.9 ± 0.97%), the representativeness of *Bacteroidetes* and *Cyanobacteria* was found to be more variable across the different DBs. The general similar distribution can be explained by the fact that the three DBs used in this study were derived from the same metagenome. Indeed, by using distinct data sources (metagenomes and different public repositories), contrasting distributions can be anticipated, as it was recently demonstrated [18]. In our study, *Alphaproteobacteria* were found to be the most represented class (72.9 ± 1.9%) followed by *Gammaproteobacteria* (18.2 ± 2.0%), *Flavobacteriia* (4.1 ± 0.5%) and unclassified *Cyanobacteria* (3.0 ± 0.7%) (Table 2). The dominance of *Alpha-* and *Gammaproteobacteria* was often reported in other marine metaproteomic studies [5, 7, 8] due to their high distribution in most marine sampling sites. Other studies focusing on sea surface sample also supported the presence of *Cyanobacteria* [47] and *Flavobacteriia* [49].

**Table 2.** Comparison of the distribution of proteins assigned at phylum and class levels for each methodology. Values represent the proportion of proteins with identical taxonomy on total identified protein using TAX-DB, NAM-DB, AM-DB or Comb-DB in both gel-free and gel-based approaches. The number of peptides detected for each protein was used as quantitative value. Taxa displaying a proportion < 1 % were gathered into “Other” category.

|  | Gel-based |  |  |  | Gel-free |  |  |  |
| --- | --- | --- | --- | --- | --- | --- | --- | --- |
|  | TAX-DB | NAM-DB | AM-DB | Comb-DB | TAX-DB | NAM-DB | AM-DB | Comb-DB |
| **Phylum** |  |  |  |  |  |  |  |  |
| *Proteobacteria* | 87.2 | 92.7 | 95.2 | 89.8 | 87.6 | 92.9 | 91.5 | 90.0 |
| *Cyanobacteria* | 6.3 | 2.9 | 2.0 | 4.0 | 2.6 | 4.8 | 1.6 | 2.0 |
| *Bacteroidetes* | 4.9 | 4.2 | 2.4 | 5.0 | 7.6 | 1.4 | 6.1 | 5.8 |
| Other (<1%) | 1.6 | 0.2 | 0.4 | 1.2 | 2.2 | 0.9 | 0.8 | 2.2 |
| **Class** |  |  |  |  |  |  |  |  |
| *Alphaproteobacteria* | 73.7 | 81.6 | 79.9 | 74.3 | 69.7 | 68.4 | 68.4 | 67.5 |
| *Gammaproteobacteria* | 12.9 | 11.5 | 14.8 | 14.4 | 18.0 | 25.7 | 24.4 | 23.5 |
| *Flavobacteriia* | 6.5 | 3.7 | 2.2 | 3.9 | 6.3 | 3.5 | 4.9 | 4.8 |
| *Unclassified Cyanobacteria* | 3.9 | 2.7 | 1.8 | 5.3 | 2.5 | 1.4 | 1.2 | 2.1 |
| Other (<1%) | 3.0 | 0.5 | 1.2 | 2.1 | 3.6 | 1.0 | 1.0 | 2.2 |

At the order level and below, the choice of DB was found to affect both qualitatively and quantitatively the taxonomic distribution, independently of the protein fractionation approach (Figure 2, Additional files 2 and 3). Although *Pelagibacterales* and *Rhodobacterales* were found to be the most dominant taxa independently of the methodology, *Pelagibacterales* were found to be overrepresented in NAM-DB and AM-DB (Figure 2a). *Pelagibacterales* are comprised of the most dominant marine microorganisms in the oceans [48] and the consistent representativeness of this order in all metaproteomes was in line with prior sea surface metaproteomic studies [5, 7, 8, 47, 53]. The observation of high protein expression profiles assigned to *Rhodobacterales* was also previously reported [50]. *Flavobacteriales* were overall more represented in the gel-free approach as well as *Cellvibrionales* but only with NAM-DB and AM-DB. *Synechococcales* were more frequently identified in the metaproteomes obtained from the gel-based approach. TAX-DB led to the characterization of many proteins from the following taxa: *Pseudomonadales*, *Rhizobiales* and *Sphingomonadales*. These taxa were either absent or rarely represented in NAM-DB or AM-DB. As stated above, TAX-DB provided the highest number of proteins successfully annotated, explaining the more diverse distribution obtained using this DB. Interestingly, the taxonomic distributions obtained with Comb-DB were found to be a good compromise between TAX-DB, NAM-DB and AM-DB (Figure 2a). Striking discrepancies in taxonomic diversity was observed between metaproteomes. As shown in the Venn diagrams provided in Figure 2b, only one quarter out of the 34 and 41 unique orders observed in gel-based and gel-free approaches respectively, was common to all DBs. Around 40 and 30% of unique orders were exclusively characterized in TAX-DB and Comb-DB in gel-based and gel-free approaches respectively, demonstrating the great performance of those DBs at extracting the broadest diversity.

**Figure 2.**
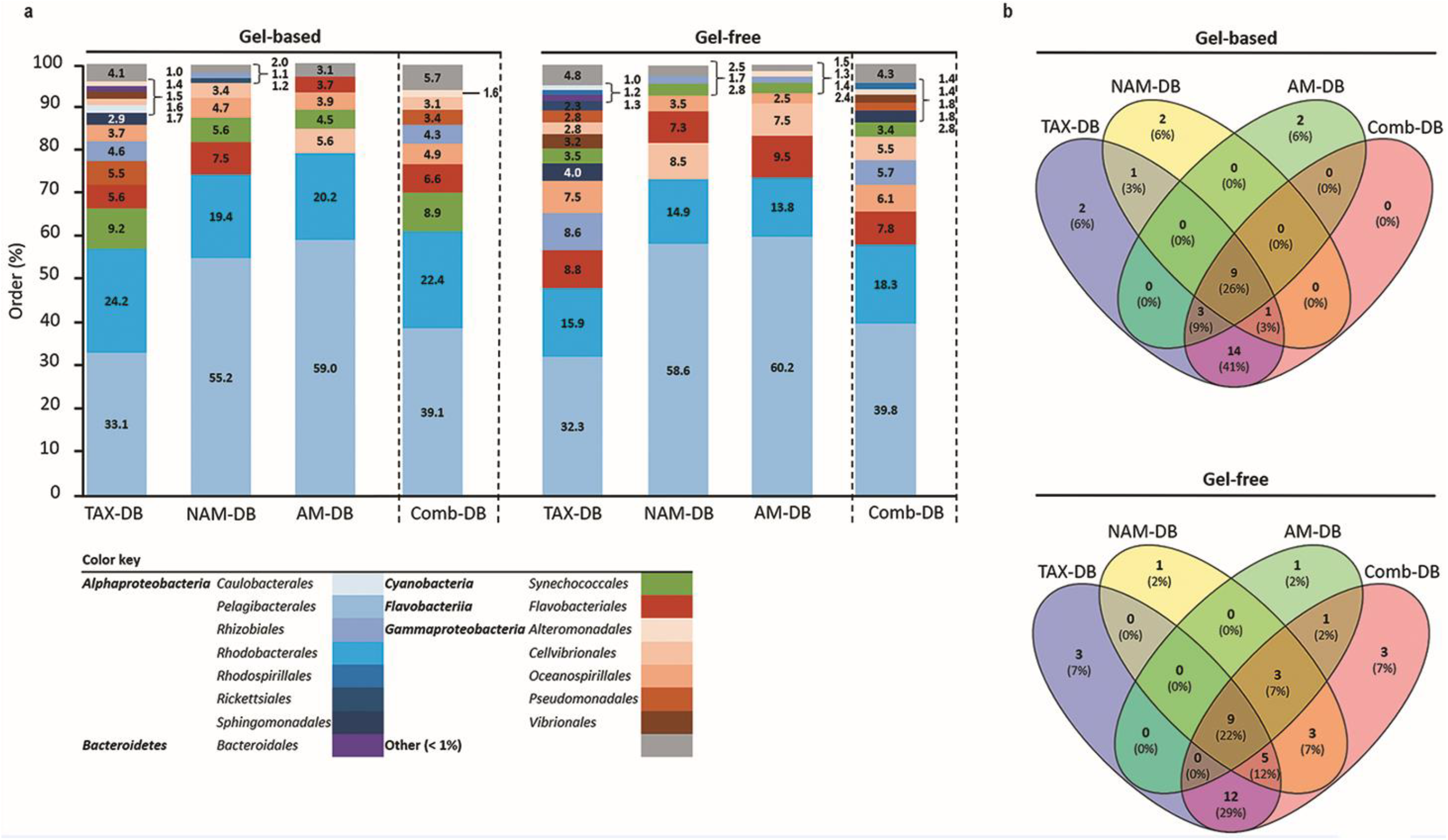
(a) Relative taxonomic composition at order level for each methodology. Values represent the proportion of proteins with identical taxonomy on total identified protein using TAX-DB, NAM-DB, AM-DB or Comb-DB in both gel-free and gel-based approaches. The number of peptides detected for each protein was used as quantitative value. Taxa displaying a proportion < 1 % were gathered into “Other” category. (b) Venn diagrams showing the number of common and unique taxa identified at order level.

### Proteomics workflow and protein search DB affect functional identification

The total number of proteins, for which a functional consensus annotation was found, decreased with the following order: TAX-DB (gel-based: 66 %, gel-free: 77 %) > AM-DB (gel-based: 61 %, gel-free: 54 %) > NAM-DB (gel-based: 50 %, gel-free: 54 %) (Figure 1, Table S2). Using Comb-DBs, 59 and 67% of functional annotation were observed in gel-based and gel-free approach respectively. In all metaproteomes, the 60 kDa chaperonin was found to be the most abundant protein (Figure 3a). The prevalence of chaperonin proteins was previously observed in other marine metaproteomic studies [7, 47, 53]. The 60 kDa chaperonin is an essential protein involved in large range of protein folding and could potentially act as signaling molecule [54]. Moreover, this protein is found in nearly all bacteria. Some taxa, such as *Alphaproteobacteria* or *Cyanobacteria*, often contain several 60 kDa chaperonin homologs [55]. On top of its ubiquity and its vital role, the abundance of the 60 kDa chaperonin could be interpreted as a response to environmental stresses exposure [7, 47, 53].

**Figure 3.**
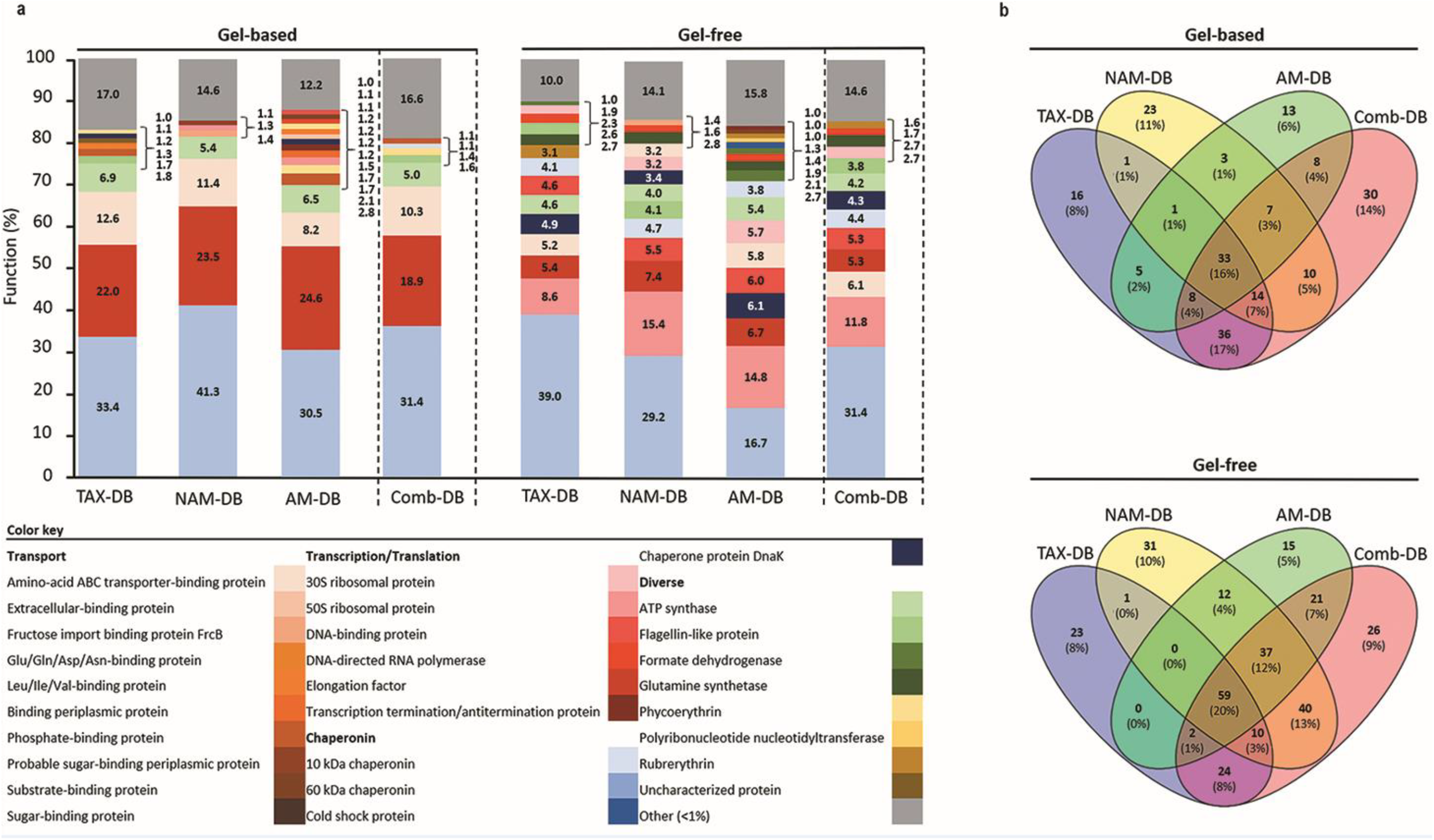
(a) Relative functional composition for each methodology. Values represent the proportion of proteins with identical functional name on total identified protein using TAX-DB, NAM-DB, AM-DB or Comb-DB in both gel-free and gel-based approaches. The number of peptides detected for each protein was used as quantitative value. Functional name displaying a proportion < 1 % were gathered into “Other” category. (b) Venn diagrams showing the number of common and unique protein functions.

Protein fractionation (gel-based *versus* gel-free) was found to affect both qualitatively and quantitatively the functional distribution as shown in Figure 3. The gel-free approach provided the greatest diversity of protein functions in comparison to the gel-based approach (Figure 3a). Only 16 and 20% of the protein functions were found to be common in all DBs from the gel-based and gel-free approaches respectively (Figure 3b). In the gel-based approach, three main functions namely the elongation factor protein, the amino-acid ABC transporter-binding protein, and the ATP synthase were observed in all DBs (Figure 3a). In contrast, in the gel-free approach, a higher number of abundant proteins was observed, including: 50S ribosomal proteins, elongation factor protein, ATP synthase, DNA-binding protein, amino-acid ABC transporter-binding protein, 10 kDa chaperonin and the chaperone protein DnaK (Figure 3a). In both proteomics approaches, each individual DB allowed the characterization of a significant number of unique protein functions (Figure 3b). Comb-DB proved to be effective at merging the results obtained from each individual DB, leading to the highest number of identified functions.

### Metaproteomic workflow alters biological interpretation

All proteins accurately annotated at both taxonomic (order rank) and functional levels were clustered and visualized into heatmaps for each DB (Figures 4). Interestingly, in 5 out of 6 heatmaps derived from NAM-DB, AM-DB and TAX-DB, *Pelagibacterales* was found to be a taxonomic cluster that stood out from all other taxa comprising of *Rhodobacterales*, *Rhizobiales, Pseudomonadales, Oceanospirillales, Cellvibrionales*, *Flavobacteriales* or *Synechococcales*. An exception was observed for TAX-DB in the gel-based approach where *Rhodobacterales* formed a distinct cluster instead of *Pelagibacterales*. Both *Pelagibacterales* and *Rhodobacterales* clustered apart together from all other taxa when using the Comb-DB. Regarding the functional clustering, the 60 kDa chaperonin was found to stand out all other functions apart from NAM-DB in the gel-based approach. Despite the similar trend observed for the most abundant taxa and most represented protein functions for all metaproteomes, Figure 4 clearly shows that the methodology was found to significantly alter the structure/function network.

**Figure 4.**
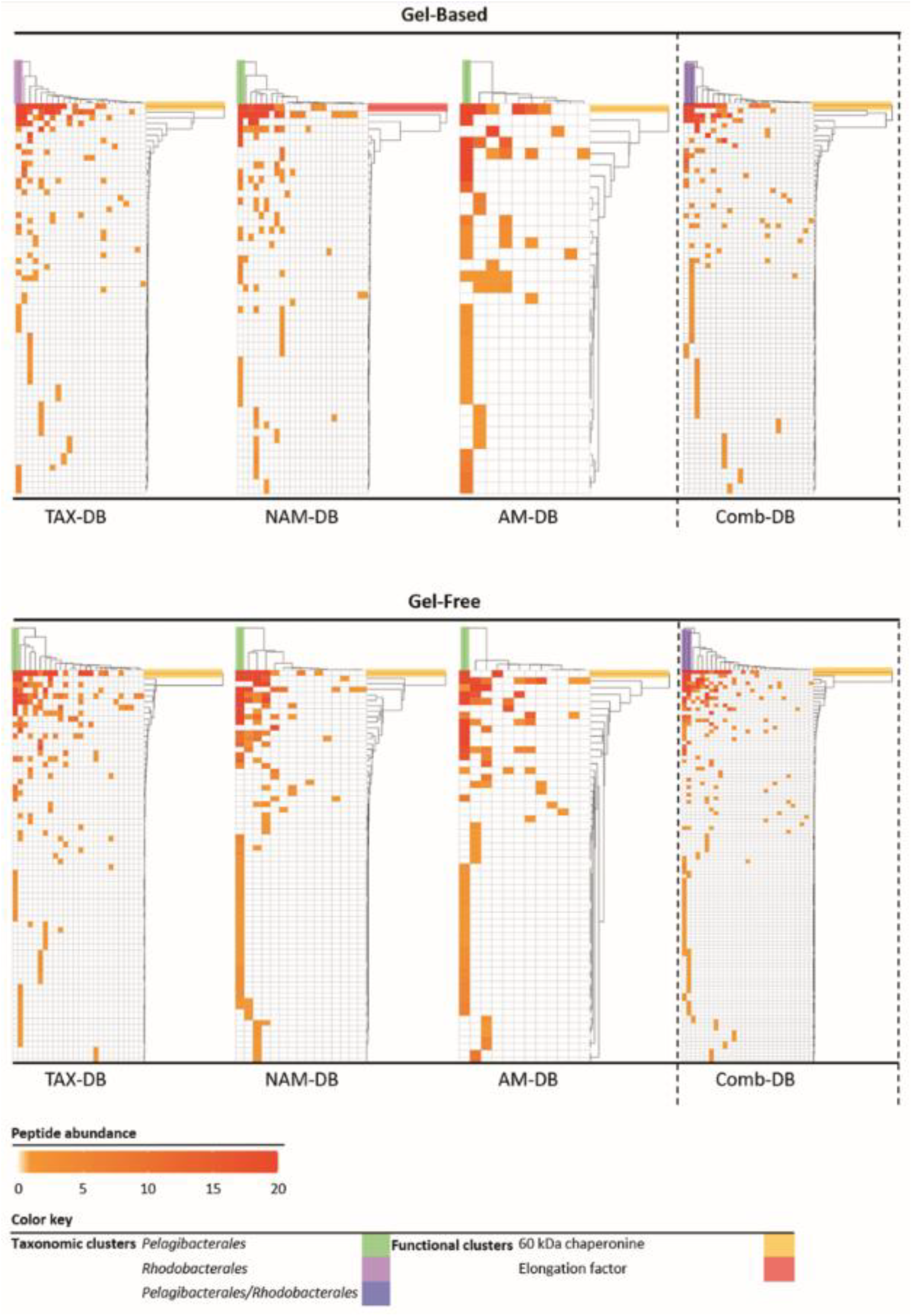
Heatmaps of the taxonomic (top clusters) and the functional (right clusters) linkages for each methodology. Proteins annotated at both order and functional levels were ranked according to the number of identified peptides. Clusters were determined using complete linkage hierarchical clustering and Euclidean distance metric.

Interestingly, the detection in all metaproteomes of numerous transporters with a broad range of substrates across different taxa could be interpreted as a strategy allowing bacteria to survive under nutrient-limited environments (Figure 5) [56, 57]. Proteins involved in amino-acid/peptide and carbohydrate transport were the most abundant transporters. In contrast, other proteins crucial in environmental processes, such as phosphorous, iron or vitamin transporters were heterogeneously characterized in a limited number of metaproteome, highlighting the risk of building upon incomplete visualization of community members coping with oligotrophic conditions.

**Figure 5.**
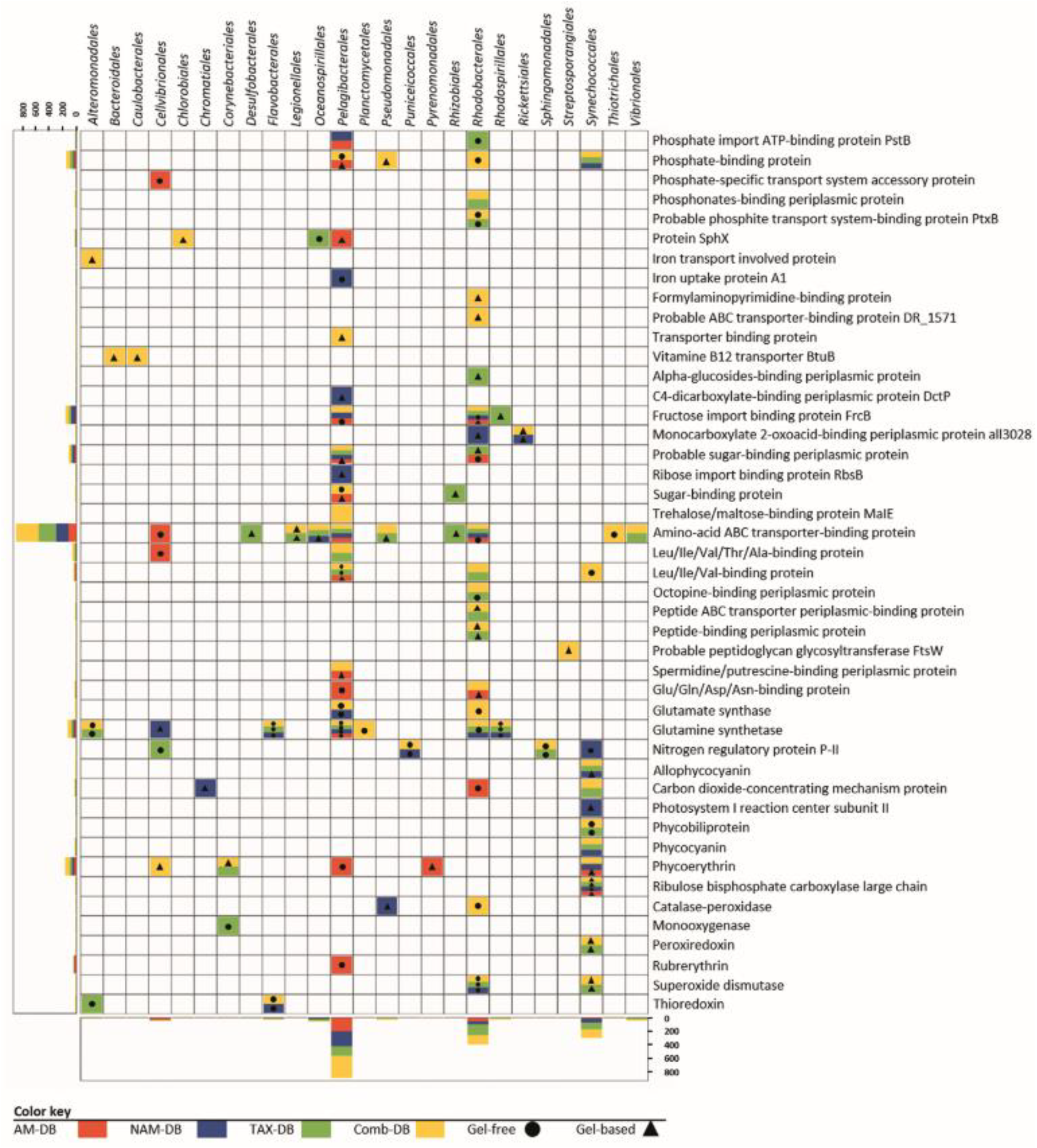
Diversity and taxonomic distribution of proteins involved in nutrient transport, nitrogen assimilation, light harvesting and oxidative stress response for each methodology. Horizontal and vertical bar charts correspond to the total number of peptides detected for a given function (y axis) or order (x axis) in all metaproteomes. The lack of symbol in colored boxes means that the protein was observed in both gel-free and gel-based approaches.

The detection of amino-acid/peptide transporters, together with other proteins involved in nitrogen metabolism (i.e. glutamate and glutamine synthase or nitrogen regulatory protein PII), strengthened the nitrogen-depleted habitat hypothesis and suggested that a wide range of bacteria were metabolically adapted to survive under nitrogen depletion [58, 59]. Surprisingly, only 3 of the 14 proteins involved in nitrogen metabolism were observed in the gel-based approach, emphasizing how protein fractionation could alter the understanding of nitrogen cycle.

The detection of proteins involved in light-harvesting, photosynthesis and oxidative stress response was found to be particularly dependent of the workflow (Figure 5). A total of 16 of the 26 proteins were characterized in only one metaproteome, showing that a robust experimental design using multiple methodologies will improve the understanding of the microbial light response. Indeed, combining the information found in all metaproteomes helped at depicting the variety of pigments belonging to photoautotrophs or photoheterotrophs [60]. The characterization of the carbon dioxide-concentrating mechanism protein Ccmk together with the ribulose bisphosphate carboxylase (RuBisCO) informed on how primary producers, such as *Synechococcales* and *Rhodobacterales* overcome inorganic carbon limitation [47, 61]. Overall, several oxidative stress-related proteins and numerous chaperonin proteins were identified in all metaproteomes, suggesting the adaptability of the microbial community to cope with oxidative stress. As a reminder, surface water samples were collected in summer at the surface of the Mediterranean Sea, where high solar irradiance was encountered. Chaperones are essential for coping with UV-induced protein damage and maintaining proper protein function [62]. Consequently, those metaproteomics results suggest that strategies used by microorganisms to cope with high solar radiation could be similar to the ones extensively described in axenic cultures using microcosms experiments [62, 63].

## Conclusion

Metaproteomics enables to progress beyond a mere descriptive analysis of microbial community diversity and structure, providing specific details on which bacteria, and which pathways of those key players, are impacted by possible perturbations. Nevertheless, using this powerful tool without fully apprehending the limitations could lead to significant misinterpretations, especially in the case of comparative metaproteomic studies. This study clearly evidenced the implications of critical decisions in metaproteomic workflow. Our findings lead to the general recommendation of diversifying when possible the protein search database as well as protein fractionation, especially if only one condition/ecosystem was studied. A robust diversified workflow allows crossing information from multiple metaproteomes in order to accurately describe the functioning of microbial communities. In a comparative metaproteomic study however, the best compromise relies on the creation of a combined DB. Our findings will undoubtedly serve future studies aiming at reliably capturing how microorganisms operate in their environment.

## Acknowledgements

This work was supported by the Royal Society, UK (RG160594), the “Belgian Fund for Scientific Research (Grand equipment - F.R.S - FNRS) and the Federal Ministry of Education and Research (BMBF, grant no 031 A535A). The funders had no role in study design, data collection and analysis, decision to publish, or preparation of the manuscript. Augustin Géron is the recipient of a 50/50 match funding scholarship between the University of Stirling (Scotland, UK) and the University of Mons (Belgium).The authors acknowledge the use of de.NBI cloud and the support by the High Performance and Cloud Computing Group at the Zentrum für Datenverarbeitung of the University of Tübingen. We want to personally acknowledge Jules Kerssemakers (German Cancer Research Center), Manuel Prinz and Katrin Leinweber (Technische Informationsbibliothek, TIB.eu) for code review, critical thoughts, and software publication advice.

## Author contributions

SMS conceived the study, performed water sampling, protein extraction and mass spectrometry analysis. AG, JW and SMS participated in the design of the mPies program. JW developed the mPies program. AG analyzed all data and wrote the manuscript. AG and JW prepared the figures. SMS, RW and PL contributed resources. All authors edited the manuscript and approved the final draft.

