## Additional file1 for "Deciphering the functioning of microbial communities: shedding light on the critical steps in metaproteomics"

**Additional file 1:** Taxonomic and functional protein annotation summary. Comparison of the proportion of proteins for which a consensus taxonomic or functional annotation was found for each methodology. Values represent the percentage of annotated proteins on total identified protein using TAX-DB, NAM-DB, AM-DB or Comb-DB in both gel-free and gel-based approaches.

|  |  | **Gel-based** |  |  |  | **Gel-free** |  |  |
| --- | --- | --- | --- | --- | --- | --- | --- | --- |
| **Taxonomic level** | **TAX-DB** | **NAM-DB** | **AM-DB** | **Comb-DB** | **TAX-DB** | **NAM-DB** | **AM-DB** | **Comb-DB** |
| Domain | 100 | 97 | 96 | 100 | 99 | 94 | 93 | 99 |
| Phylum | 95 | 92 | 91 | 94 | 95 | 87 | 88 | 92 |
| Class | 81 | 78 | 81 | 80 | 85 | 77 | 80 | 81 |
| Order | 70 | 41 | 45 | 51 | 71 | 42 | 44 | 52 |
| Family | 64 | 34 | 37 | 44 | 66 | 32 | 33 | 43 |
| Genus | 50 | 7 | 13 | 28 | 43 | 8 | 9 | 16 |
| Function | 66 | 50 | 61 | 59 | 77 | 54 | 54 | 67 |
