## Additional file4 for "Deciphering the functioning of microbial communities: shedding light on the critical steps in metaproteomics"

**Additional file 4:** mPies program features

- Project name: mPies
- Project home page: <https://github.com/johanneswerner/mPies>
- Archived version: 10.5281/zenodo.2539710
- Operating system(s): Linux
- Programming language: Python 3
- Other requirements: Conda, other packages installed with mPies: CD-Hit AuxTools (10.1093/bioinformatics/bts565), Diamond (10.1038/nmeth.3176), ETE3 (http://mbe.oxfordjournals.org/content/early/2016/03/21/molbev.msw046), FragGeneScan (10.1093/nar/gkq747), MEGAHIT (10.1093/bioinformatics/btv033), Prodigal (10.1186/1471-2105-11-119), Snakemake (10.1093/bioinformatics/bts480), Spades (10.1089/cmb.2012.0021), Trimmomatic (10.1093/bioinformatics/btu170). At least 150 GB of disk space (depending on the used database), 64 GB memory (if assembly is necessary) and 16 CPUs are recommended for running mPies.
- License: GNU GPL-3.0
- Any restrictions to use by non-academics: none
