## Additional file5 for "Deciphering the functioning of microbial communities: shedding light on the critical steps in metaproteomics"

**Additional file 5:** Examples of functional consensus annotations returned using mPies based on the 20 best blast hits. The three examples were retrieved using second-round search NAM-DB in gel-free approach.

**Example A:** automatic mPies annotation

| **Candidate annotations from the top alignment hits** | **Frequency** | **Similarity (%)** |
| --- | --- | --- |
| Glutamine synthetase | 16 | 80 |
| Glutamine synthetase nodule isozyme | 1 | 5 |
| Glutamine synthetase root isozyme 3 | 1 | 5 |
| Glutamine synthetase root isozyme A | 1 | 5 |
| Glutamine synthetase root isozyme B | 1 | 5 |
| **Consensus functional annotation** |  |  |
| Glutamine synthetase |  |  |

**Example B: manual annotation**

| **Candidate annotations from the top alignment hits** | **Occurrences** | **Proportion (%)** |
| --- | --- | --- |
| 60 kDa chaperonin 1 | 9 | 45 |
| 60 kDa chaperonin 2 | 3 | 15 |
| 60 kDa chaperonin 3 | 3 | 15 |
| 60 kDa chaperonin | 2 | 10 |
| 60 kDa chaperonin 4 | 1 | 5 |
| 60 kDa chaperonin 5 | 1 | 5 |
| 60 kDa chaperonin 7 | 1 | 5 |
| **Consensus functional annotation** |  |  |
| 60 kDa chaperonin |  |  |

**Example C: no consensus annotation**

| **Candidate annotations from the top alignment hits** | **Occurrences** | **Proportion (%)** |
| --- | --- | --- |
| 3-hexulose-6-phosphate synthase | 2 | 25 |
| 5,6,7,8-tetrahydromethanopterin hydro-lyase | 2 | 25 |
| 5,6,7,8-tetrahydromethanopterin hydro-lyase | 2 | 25 |
| Bifunctional enzyme Fae/Hps | 2 | 25 |
| **Consensus functional annotation** |  |  |
| NA |  |  |
